# Fast-spiking neurons in monkey orbitofrontal cortex underlie economic value computation

**DOI:** 10.1101/2025.04.06.647503

**Authors:** Tomoaki Murakawa, Takashi Kawai, Yuri Imaizumi, Hiroshi Yamada

**Affiliations:** Academic service office for the medical science area, University of Tsukuba, 1-1-1 Tenno-dai, Tsukuba, Ibaraki 305-8577, Japan; The Picower Institute for Learning and Memory, Department of Biology and Department of Brain and Cognitive Sciences, Massachusetts Institute of Technology, Cambridge, MA 02139, USA; College of Medical Sciences, University of Tsukuba, 1-1-1 Tenno-dai, Tsukuba, Ibaraki 305-8577, Japan; Division of Biomedical Science, Institute of Medicine, University of Tsukuba, 1-1-1 Tenno-dai, Tsukuba, Ibaraki 305-8577, Japan

**Keywords:** Orbitofrontal cortex, inhibitory interneuron, monkey, economic behavior

## Abstract

Inhibitory interneurons are fundamental constituents of cortical circuits that process information to shape economic behaviors. However, the role of inhibitory interneurons in this process in the core cortical reward region, the orbitofrontal cortex (OFC), remains elusive. Here, we show that presumed parvalbumin-containing GABAergic interneurons (fast-spiking neurons, FSNs) differ from regular-spiking pyramidal neurons (RSNs) in computing economic value. While monkeys perceived a visual lottery for probability and magnitude of rewards, identified FSNs constituted 12% of OFC neurons and showed high-frequency firing rates and wide dynamic ranges, which are both key intrinsic cellular characteristics regulating cortical computation. We found that FSNs showed stronger neural modulation compared to RSNs in signaling probability and magnitude information. Unambiguously, both neural populations signaled the expected values, but FSNs processed the reward information more strongly under the influence of dynamic range. Thus, stronger FSN signals may appear with wider dynamic range for expected value computation.

## INTRODUCTION

Functions of circuits that regulate economic behaviors rely on the ability of interneurons to regulate information flow in the cortical and subcortical structures (Chamberland et al., 2023; Giordano et al., 2023; Gonzalez-Burgos et al., 2005; Wilbers et al., 2023; Yamada et al., 2016). This computational process is thought to depend on circuit structures that regulate the economic behavior of animals. Indeed, cortical inhibitory dysfunction results in various diseases, including mental disorders (Allami et al., 2025; Hattori et al., 2017). Since excitatory neurons constitute the majority of neurons at one of the core cortical reward centers, the orbitofrontal cortex (OFC), they have been extensively examined in relation to economic behavior to obtain rewards (Chen et al., 2023; Imaizumi et al., 2022; Padoa-Schioppa, 2013; Pastor-Bernier et al., 2021; Rich and Wallis, 2016; Yamada et al., 2021; Yamada et al., 2018). However, it remains unclear how OFC inhibitory interneurons are involved in shaping economic behaviors, especially in macaque monkeys, close relatives to humans.

Parvalbumin-containing GABAergic interneurons have generally been identified as fast-spiking neurons (FSNs) in the brain based on their narrow spike waveforms (Cauli et al., 1997; DeFelipe, 1997; Kawaguchi and Kondo, 2002; Kawaguchi et al., 1995). Although cortical excitatory activity is regulated by inhibitory interneurons (Bartho et al., 2004; Giordano et al., 2023; Hatch et al., 2017; Owen et al., 2018; Wilbers et al., 2023), FSN activity in cortical regions, particularly in monkeys, has been examined in only a few studies of cognitive and motor task performance (Constantinidis and Goldman-Rakic, 2002; Gonzalez-Burgos et al., 2005; Kawai et al., 2019; Mitchell et al., 2007; Povysheva et al., 2006; Wang et al., 2016). To the best of our knowledge, few studies have examined the role of FSNs in the OFC during economic behavior in both monkeys and rodents. This is largely because FSNs constitute a minority of neurons; therefore, only limited data can be obtained from a single study. Given this limitation, it is challenging to elucidate the inhibitory mechanisms of FSNs in monkey OFCs which process economic- value computations from the inhibitory dysfunction as suspected (Hattori et al., 2017; Prevot and Sibille, 2021).

In the present study, we aimed to understand how FSNs regulate OFC activity during gambling behavior in monkeys. We differentiated FSNs from presumed regular-spiking pyramidal neurons (RSNs) based on extracellularly recorded spike waveforms in the OFC of behaving monkeys. We addressed two critical issues in examining the role of FSNs: 1) How are FSNs in the OFC of behaving monkeys involved in perceiving expected values, that is, probability multiplied by the magnitude of the reward? 2) How does the activity of FSNs differ from that of RSNs in the OFC when computing expected values? Our results suggest that FSNs may compute expected values in coordination with RSNs in the OFC, with this process influenced by the dynamic range, a key factor encoding information (Laughlin, 1981).

## RESULTS

### Identification of FSNs and their basic firing properties

We studied a total of 377 neurons in the OFC of behaving monkeys during a single-cue task (Figure 1A). We previously reported monkey behavior during a choice task (Yamada et al., 2021) (Figure 1B-D). In short, monkeys chose the option with the higher expected value (i.e., probability times magnitude). While the monkeys viewed a visual lottery composed of green (reward magnitude) and blue (reward probability) pie charts (Figure 1A), neuronal activity was recorded in the OFC (Figure 1E; medial [mOFC], 14O; central [cOFC], 13M). We classified the neurons into FSNs and RSNs based on spike waveforms, following procedures previously used in rat and monkey studies (Bartho et al., 2004; Yamada et al., 2016).

**Figure 1.**
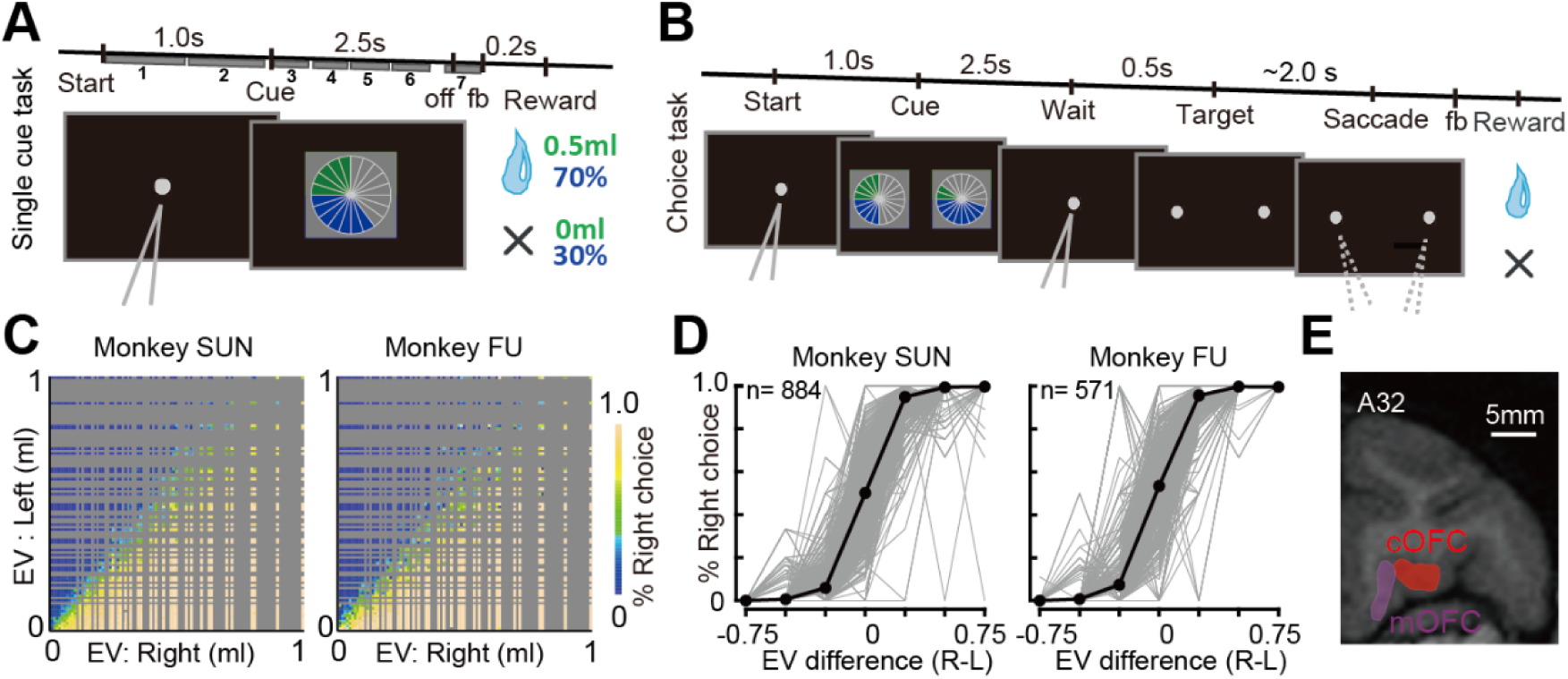
Task, behavior, and recording sites. **A.** Sequence of events during the single-cue task. A single pie chart with green and blue segments was presented visually to the monkeys. The magnitude and probability were indicated by green and blue segments, respectively. Seven 0.5-s analysis time periods were represented by gray boxes numbered from 1 to 7 (1: Start1, 2: Start2, 3: Cue1, 4: Cue2, 5: Cue3, 6: Cue4, and 7: Pre-fb). The feedback sound (fb) was followed by reward delivery. **B.** Choice task. Two pie charts were presented visually to the monkeys on the left and right sides of the center. After visual fixation on the central point, it disappeared, and the monkeys chose one of the targets by fixating on it. A block of choice trials was sometimes interleaved between single-cue trial blocks. During the choice trials, neural activity was not recorded. **C.** Percentages of right target choices during the choice task plotted against the expected values (EVs, probability × magnitude) of the left and right target options. The aggregated choice data were analyzed. Gray indicates that no samples exist for such lottery combinations. **D.** Percentage of correct target choices estimated in each recording session (gray lines) plotted against the difference in expected values (right minus left). The choice data were segmented into seven conditions based on the difference in expected values: -1.0 ∼ -0.5, -0.5 ∼ -0.3, -0.3 ∼ -0.1, -0.1 ∼ 0.1, 0.1 ∼ 0.3, 0.3 ∼ 0.5, and 0.5 ∼1.0. The black plots indicate the mean values. The total number of recording sessions for choice trials are shown. **E.** Illustration of neural recording areas based on coronal MRI. Neurons were recorded from the medial (mOFC, 14O, orbital part of area 14) and central parts of the orbitofrontal cortex (cOFC, 13M, medial part of area 13) at the A31–A34 anterior-posterior (A-P) level. These figures were taken from Yamada et al. (2021) and slightly modified in A and B.

To classify the FSNs, we identified two critical characteristics of the spike waveform (Figure 2A): peak width (i.e., width at half maximum of the negative peak amplitude; see green inset in Figure 2A) and peak-to-valley width (i.e., time from negative peak to valley) (see Materials and Methods). A scatter plot of peak width against peak-to-valley width for all neurons revealed the sharpest waveform neurons, mainly differentiated by peak-to-valley width (Figure 2A). We classified FSNs as neurons within one cluster that exhibited the narrowest spike waveforms (Figure 2A, green; insets). Other neurons were classified as RSNs according to the previous study by Bartho et al., 2004. The identified FSNs accounted for approximately 12% of the recorded OFC neurons (42/377; cOFC, n = 25; mOFC, n = 17). We previously reported the activity of RSNs (Chen et al., 2023; Chen et al., 2025; Imaizumi et al., 2022; Yamada et al., 2021), but not that of FSNs in this dataset during the cued lottery task. We note that we did not record OFC activity during the choice task.

**Figure 2.**
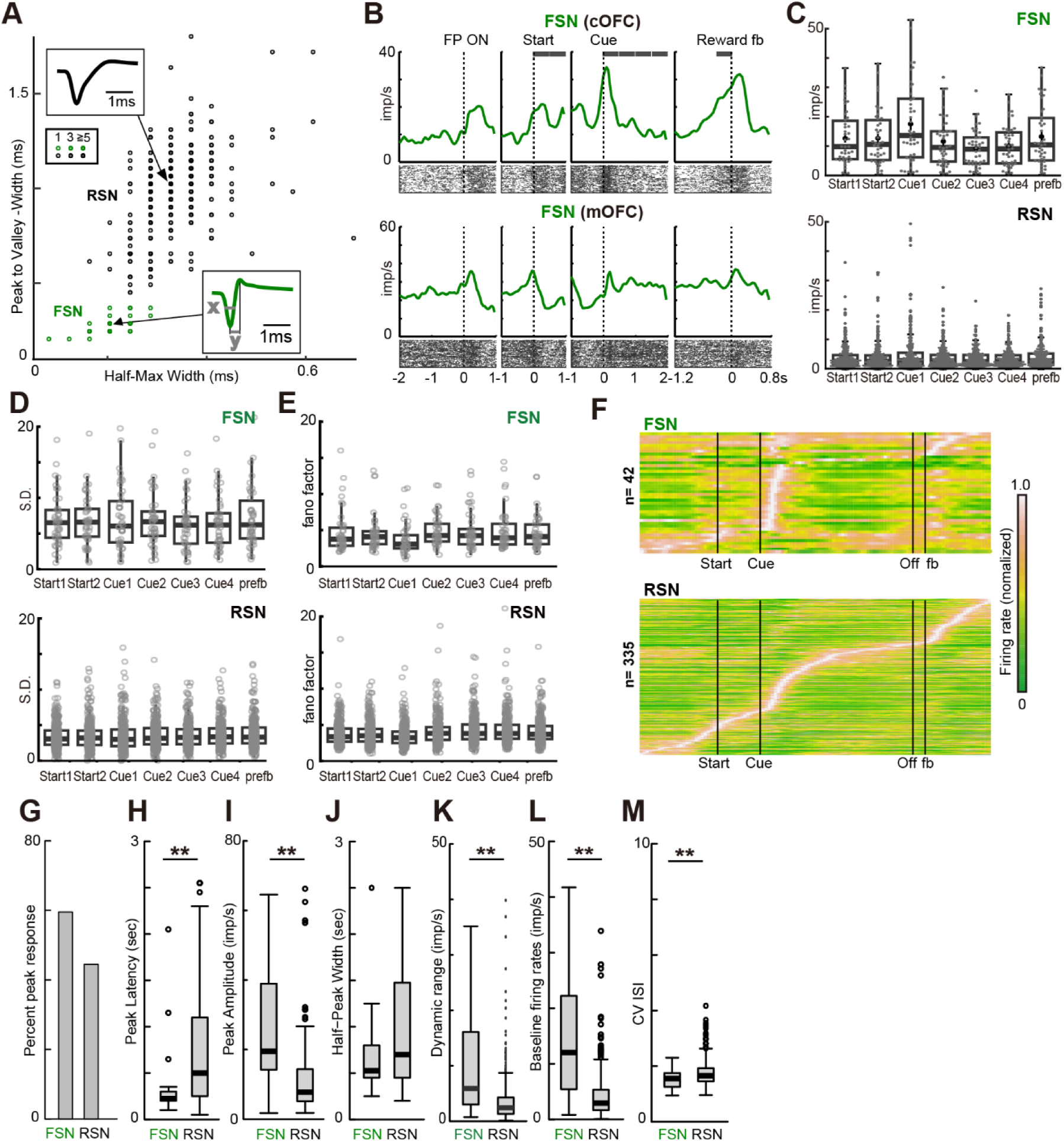
Classification of FSNs and their basic activity properties during task trials. **A.** Scatter plots of mean spike waveform duration (x: width at half maximum of the negative peak amplitude; y: width from peak to valley; inset) for OFC neurons. FSNs are defined as neurons in a cluster that exhibit narrow spike waveforms (green) in line with Bartho et al 2004. Neurons in clusters with wider spike waveforms were classified as RSNs. Darkness in the overlaid scatter plot indicates the number of neurons. **B.** Two examples of FSN activity recorded from the cOFC of monkey SUN and the mOFC of monkey FU during a single-cue task. Rasters and histograms were aligned for each behavioral event. The seven gray bars at the top indicate the 0.5-s analysis periods. All histograms (50-ms bins) were smoothed using a Gaussian kernel (50 ms). **C.** Average firing rates of 42 FSNs and 335 RSNs during seven analysis periods. **D.** Across-trial variance estimated during seven analysis periods estimated as S.D. **E.** fano factor estimated during seven analysis periods. **F.** Color map histograms of FSN and RSN activity. Each horizontal line indicates neural activity aligned with cue onset, averaged across all lottery conditions. Neuronal firing rates were normalized to peak activity. **G.** Percentage of neurons showing peak activity during cue presentation. **H.** Peak activity latency after cue presentation. **I.** Firing rates at peak activity observed during cue presentation. **J.** Half-peak width, indicating the phasic nature of activity changes. **K.** Difference between the maximum and minimum firing rates. **L.** Box plots of baseline firing rates during the 1-s period before the presentation of the central fixation target. **M.** Coefficient of variation of inter-spike interval (CV ISI). In **H-M**, asterisks indicate statistical significance between the two neural populations (Wilcoxon rank-sum test, \**P* < 0.05, \*\**P* < 0.01).

Typical FSN activity recorded from the cOFC showed tonic firing at >10 Hz during most task periods, with a phasic increase in discharge for some task events (Figure 2B, top). Another example of FSN activity recorded from the mOFC showed a phasic increase in discharge at the start of a trial, followed by increases and decreases in activity throughout the trial (Figure 2B, bottom). We first examined firing rate changes throughout a trial, before and after the visual cue for probability and magnitude. Quantitative comparison of the average firing rates between 42 FSNs and 335 RSNs across the seven 0.5 s task periods (see Figure 1A, gray bars labeled from 1 to 7: Start1, Start2, Cue1, Cue2, Cue3, Cue4, and Pre-fb, see also gray bars in Figure 2B, top) showed higher firing rates in FSNs than in RSNs throughout a task trial (Figure 2C, two-way

ANOVA, n = 377, neuron type: F_(1,363)_ = 97.9, *P* < 0.001, task period: F_(6,363)_ = 1.94, *P* = 0.073, interaction: F_(6,363)_ = 1.01, P = 0.419). In addition, variability of spike, as i) across-trial variances (Figure 2D, two-way ANOVA, n = 377, neuron type: F_(1,363)_ = 317.8, *P* < 0.001, task period: F_(6,363)_ = 1.90, *P* = 0.077, interaction: F_(6,363)_ = 0.664, P = 0.679) and ii) fano-factor (Figure 2E, two-way ANOVA, n = 377, neuron type: F_(1,363)_ = 22.9, *P* < 0.001, task period: F_(6,363)_ = 13.6, *P* < 0.001, interaction: F_(6,363)_ = 0.657, P = 0.685), was high for FSNs compared to RSNs. Thus, the FSNs identified in the OFC of behaving monkeys showed typical characteristics of FSNs.

Specifically, FSNs changed their activity at different times during the task trials (Figure 2F). Approximately, 60% of the FSNs demonstrated peak activity during cue presentation (Figure 2G, 59.5%, 25/42), whereas a similar proportion of the RSNs showed peak activity during cue presentation (Figure 2G, 44.4%, 149/335; chi-square test: n = 377, *P* = 0.093, X^2^ = 2.82, df = 1). Peak activity with short latencies was observed in the FSNs (Figure 2H, latency, Wilcoxon rank-sum test: n = 174, *P* < 0.001, W = 2671.5, df = 1), along with higher magnitudes of activity (Figure 2I, peak firing rate: n = 174, *P* < 0.001, W = 896, df = 1). The speed of activity change was similar between the two types of neurons (Figure 2J, half-peak width, Wilcoxon rank-sum test: n = 174, P = 0.160, W = 2190.5, df = 1). In addition, the dynamic ranges (see Materials and Methods) in the FSNs were wider than those in the RSNs (Figure 2K, n = 377, *P* < 0.001, W = 263637.5, df = 1), which are critical characteristic for processing computations (Shew et al., 2009). We also confirmed that the baseline firing rates during the inter-trial interval were higher in FSNs (Figure 2M, n = 377, *P* < 0.001, W = 2794, df = 1), similar to the activity during the task trials (Figure 2C). Spike irregularity, variation of inter-spike intervals, was lower in FSNs than in RSNs (Figure 2M; n = 377, *P* = 0.007, W = 8818, df = 1). We also noted that during the feedback and reward periods, many FSNs showed increased responses relative to baseline discharge (25/42, 59.5%), in contrast to RSNs (98/335, 29.3%).

Collectively, high-frequency activity with short latency occurred in FSNs at more regular intervals, in contrast to lower firing rates at more irregular intervals in RSNs. The variability of spikes across trials was high for SFNs. Unambiguously, the dynamic range of FSNs was wider than that of RSNs, indicating that the identified FSNs exhibited characteristics that matched those of parvalbumin-containing GABAergic interneurons (Kawaguchi, 1993, 1995; Povysheva et al., 2008). We note that RSNs may be composed of multiple neuron types as suggested by the existence of multiple cortical neuronal types (Kawaguchi, 1995).

### Coding of expected values in FSNs and RSNs in raw firing rates

We examined how individual FSNs and RSNs processed information on the probability and magnitude of rewards during expected value computation. First, after cue appearance, 40–50% of the FSNs encoded the probability and magnitude of rewards until the outcome was presented (Figure 3A, left). We identified four coding types: probability, magnitude, expected value, and risk-return types (see Materials and Methods). For example, neurons signaling the expected value were found (Figure 3B), whose activity increased when either the probability or magnitude of the rewards increased (i.e., EV+ type). In addition, probability (Figure 3C, P- type) and magnitude (Figure 3D, M+ type) coding types were found for both positive and negative responses. Indeed, the neural signals carried by FSNs and RSNs were composed of a mixture of these signals (Figure 3A, left and right), including those for expected value and its components (i.e., probability and magnitude). Thus, both FSNs and RSNs signaled information for expected-value computations with a variety of neural modulation types.

**Figure 3.**
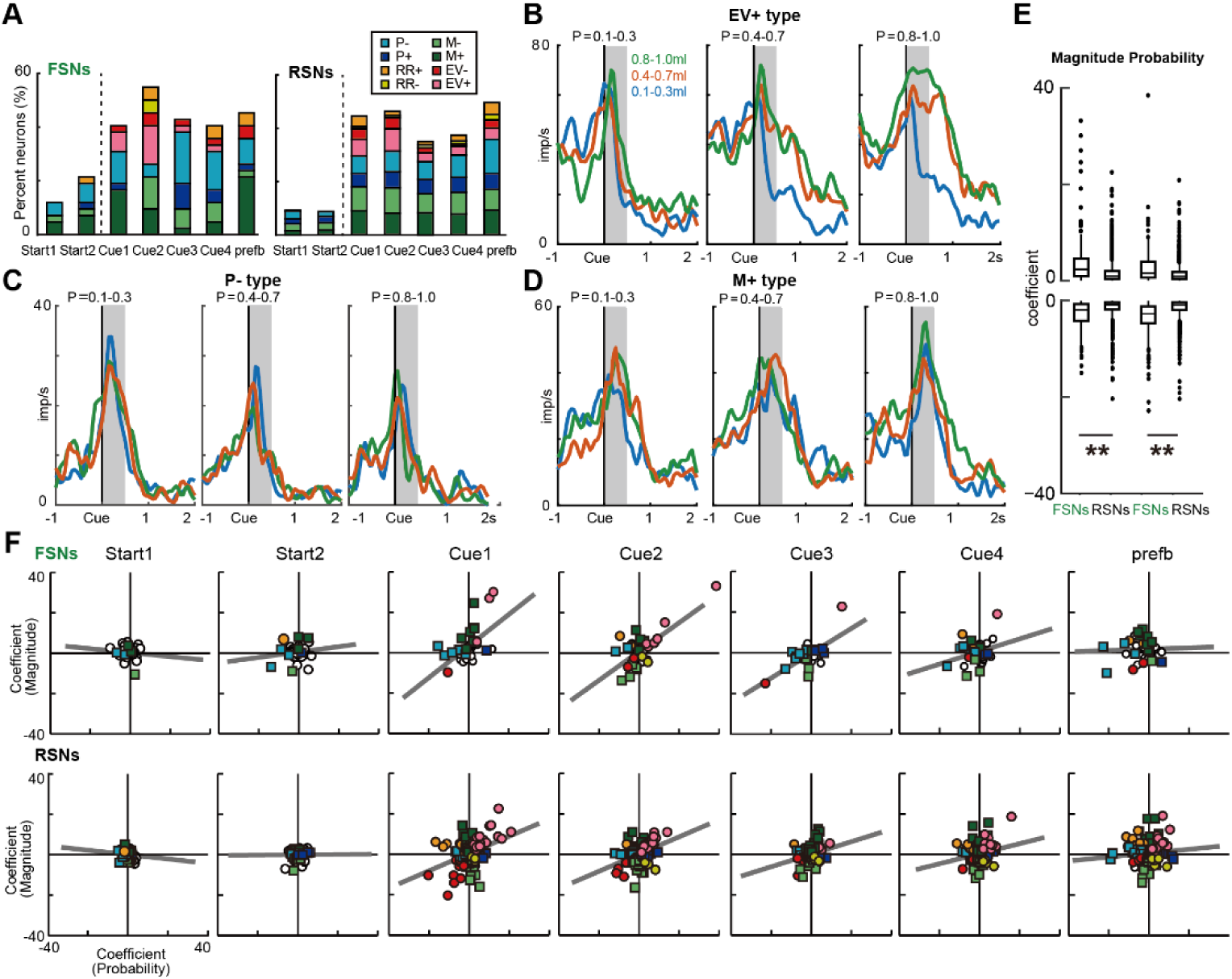
Probability and magnitude modulations of the raw firing rates in FSNs and RSNs. **A.** Percentage of neural modulation types for FSNs and RSNs during the seven analysis periods. Probability (P), magnitude (M), expected value (EV), and risk–return (RR) types were identified using the linear regression analysis. **B–D.** Examples of FSNs for EV positive regression type (EV+), P negative regression type (P-), and M positive regression type (M+) are shown. In B–D, the reward probability (P) was differentiated among low, middle, and high conditions. The reward magnitude was also differentiated among the low, middle, and high conditions using different colors. The gray-hatched time windows indicate the analysis period, Cue1. **E.** Boxplots of the regression coefficients for the probability and magnitude of rewards among positive and negative coding types. Asterisks indicate statistical significance between FSNs and RSNs (Wilcoxon rank-sum test, \**P* < 0.05, \*\**P* < 0.01). **F.** Regression coefficients for the probability and magnitude of rewards across seven analysis periods. Gray lines indicate regression slopes.

Next, we compared the neural modulations between FSNs and RSNs at the population level. Both neural populations encoded the expected values after cue presentation, as observed in regression slopes close to a 45° angle (Figure 3F, Cue1, gray line). This expected value code emerged immediately after cue appearance. Thereafter, they gradually lost these expected value signals throughout a trial, as indicated by changes in the regression slopes (Figure 3F, general linear model, n = 1885 [377 neurons ₓ 5 task periods], task period: *F* = 4.32, *P* = 0.002, df = 4). The signal change to no particular structural coding at the population level (regression slope close to 0°) occurred concurrently in both FSNs and RSNs. Regression coefficients were consistently larger in FSNs than in RSNs, and thus relatively close to a 45-degree angle throughout the trial (neuron type: *F* = 12.0, *P* < 0.001, df = 1). The same results were observed when we estimated the Pearson’s’ and Spearman’s’ correlation coefficients (see r and rho values in Tables 1 and 2); moderate correlation coefficients were observed during the Cue1 to Cue3 periods in FSNs, and smaller coefficients were observed in RSNs. These results indicate that FSNs may be closer to the expected value code than the RSNs at the population level.

**Table 1.**
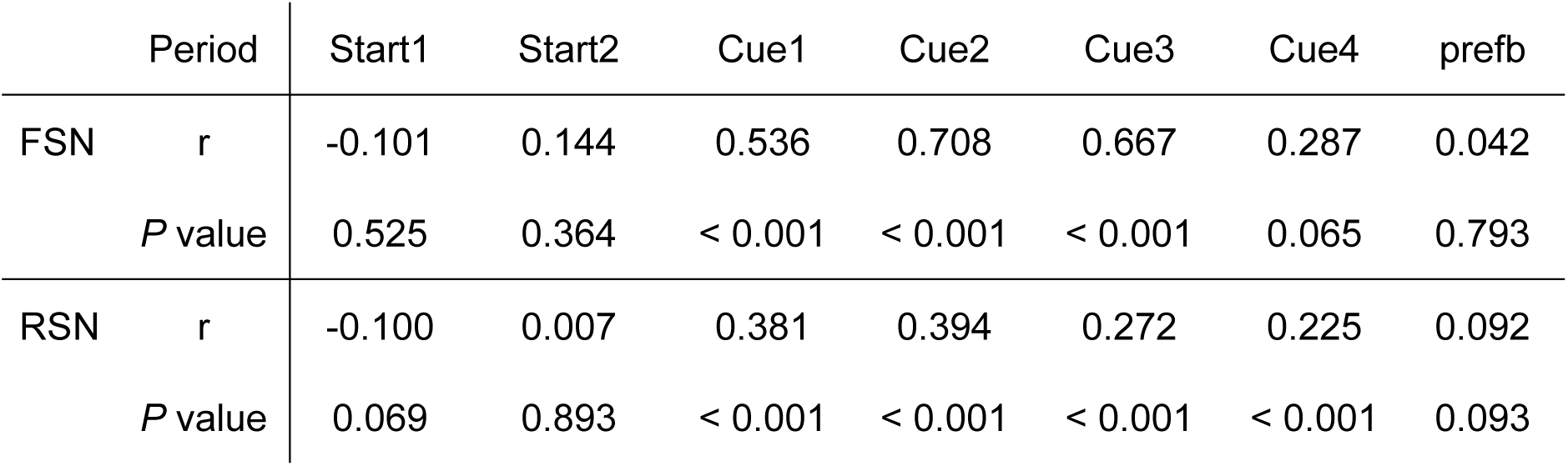
Results of Pearson’s correlation test corresponding to Figure 3F.

**Table 2.** Results of Spearman’s correlation test corresponding to Figure 3F.

| Period |  | Start1 | Start2 | Cue1 | Cue2 | Cue3 | Cue4 | prefb |
| --- | --- | --- | --- | --- | --- | --- | --- | --- |
| FSN | rho | 0.021 | 0.056 | 0.543 | 0.360 | 0.432 | 0.019 | -0.200 |
|  | P value | 0.897 | 0.725 | < 0.001 | 0.020 | 0.005 | 0.904 | 0.204 |
| RSN | rho | -0.046 | -0.034 | 0.147 | 0.280 | 0.155 | 0.085 | 0.043 |
|  | P value | 0.405 | 0.530 | 0.007 | < 0.001 | 0.005 | 0.122 | 0.434 |

Additionally, we compared the extent of neural modulation between FSNs and RSNs (Figure 3E). Neural modulation in FSNs was larger than that in RSNs, irrespective of probability or magnitude modulation (Figure 3E, four-way ANOVA, n = 3770 [377 neurons ₓ 5 task periods ₓ 2 coding types], neuron type: F_(1, 3730)_ = 229.4, *P* < 0.001; coding type: F_(1,3730)_ = 1.73, *P* = 0.189, task period: F_(4,3730)_ = 9.51, *P* < 0.001). Thus, neural modulations by FSNs (i.e., response sensitivity) were larger than those by RSNs in the raw firing rates, whereas neural modulations concurrently changed throughout a task trial at the population level.

In conclusion of the raw firing rates analysis, since the raw firing rate contains both signal and noise, these results suggested that either or both of signal and noise contribute to the higher neural modulation in FSNs (Figure 3E) due to their higher firing rates than those in RSNs (Figure 2C and L). Next, we examined this issue using z-scored firing rates for eliminating the effect of noise in the analyses.

### Coding of expected values in FSNs and RSNs using z-scored firing rates

To fairly compare information processing between FSNs and RSNs, both signals and noise must be considered to understand their contributions to firing rates. For this purpose, we normalized the firing rates to z-scores, which allowed us to directly compare the encoded information (i.e., signal contribution) between the two neural populations (Chung et al., 1987; Passaglia and Troy, 2004; Theunissen et al., 1996). We performed the same regression analyses on z-scored firing rates (see Materials and Methods; standardized regression coefficients were estimated) in comparison with the analysis results in the raw firing rates (Figure 3).

First, when analyzing the z-scored firing rates, the categorized functional types of neurons were similarly observed (Figure 4A) as the raw firing rates (Figure 3A). All neuron types were observed throughout a task trial after cue presentation. Second, changes in the extent of neural modulation throughout the task trial were similar at the population level (Figure 4B, compare with Figure 3F). Similarity in population-level codes was observed throughout the task trial (Figure 4B, compare with Figure 3F, general linear model, n = 1885 [377 neurons ₓ 5 task periods], task period: *F* = 1.7, *P* = 0.147, df = 4). The significant difference in regression slopes between neuron types was observed at the population level (general linear model, n = 1885 [377 neurons ₓ 5 task periods], neuron type: *F* = 3.93, *P* = 0.048, df = 1). This observation between FSNs and RSNs was confirmed by the correlation coefficient analyses (see Tables 3 and 4). Fourth and most importantly, the stronger neural modulation in FSNs than that in RSNs was similarly observed (Figure 4C, four-way ANOVA, n = 3770 [377 neurons ₓ 5 task periods ₓ 2 coding types], neuron type: F_(1, 3730)_ = 14.7, *P* < 0.001, coding sign: F_(1, 3730)_ = 1.25, *P* = 0.264, coding type: F_(1, 3730)_ = 1.78, *P* = 0.182, task period: F_(4, 3730)_ = 12.5, *P* < 0.001), in comparison with the raw firing rates in FSNs (Figure 3E). Thus, most analyses results were similar between the raw and z-scored firing-rate analyses.

**Figure 4.**
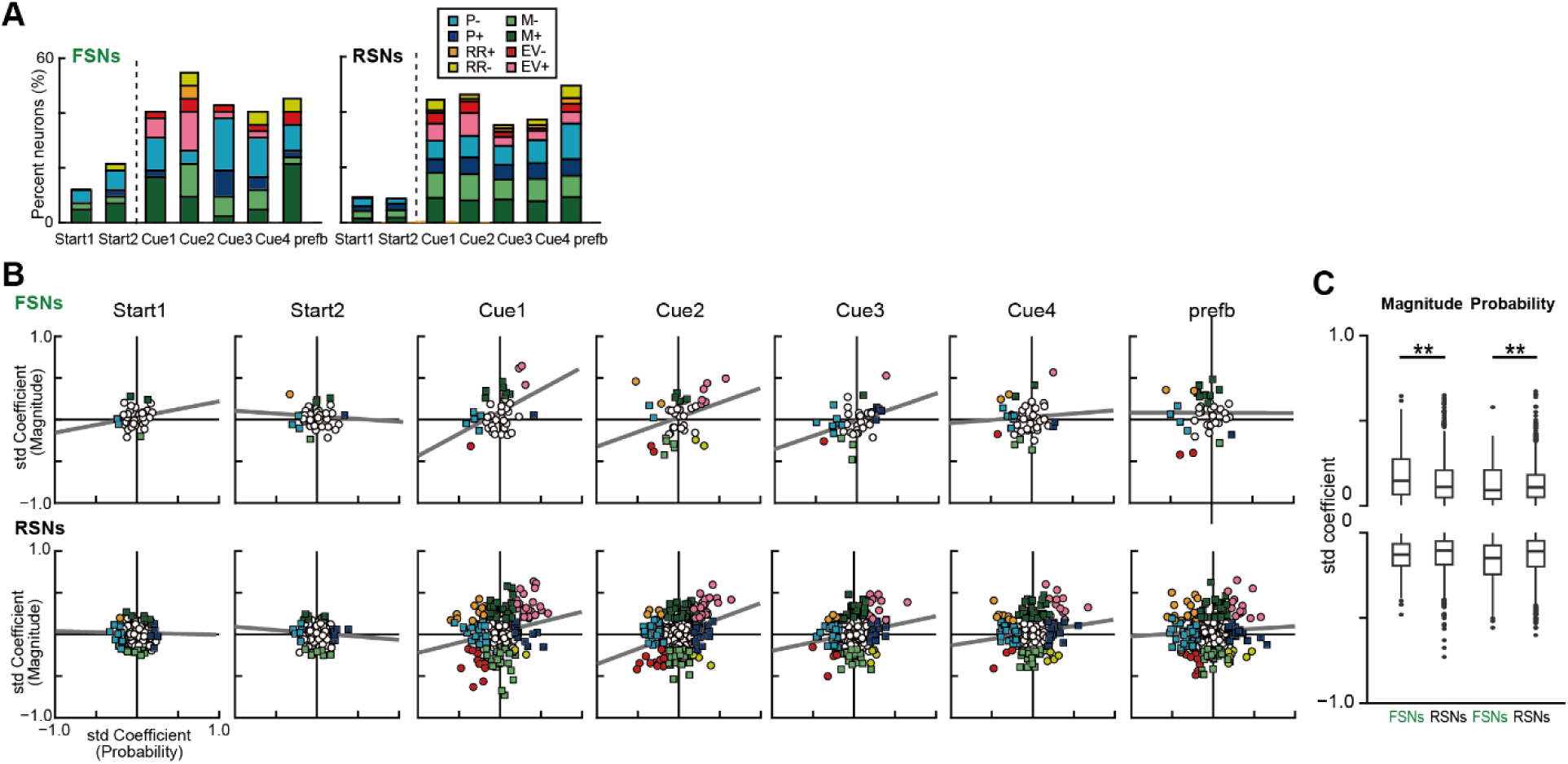
Probability and magnitude modulations of the z-scored firing rates in FSNs and RSNs. **A.** Percentage of neural modulation types for FSNs and RSNs during the seven analysis periods. Probability (P), magnitude (M), expected value (EV), and risk–return (RR) types were detected based on the linear regression analysis. **B.** Standardized (std) regression coefficients for the probability and magnitude of rewards during a task trial. The gray lines indicate regression slopes. The r and p values at the top right indicate the correlation coefficient and statistical significance, respectively. **C.** Boxplots of the standardized regression coefficients for the probability and magnitude of rewards among positive and negative coding types. Asterisks indicate statistical significance between FSNs and RSNs (Wilcoxon rank-sum test, \**P* < 0.05, \*\**P* < 0.01). n.s. indicates no significant difference between FSNs and RSNs.

**Table 3.** Results of Pearson’s correlation test corresponding to Figure 4B.

| Period |  | Start1 | Start2 | Cue1 | Cue2 | Cue3 | Cue4 | prefb |
| --- | --- | --- | --- | --- | --- | --- | --- | --- |
| FSN | Cor | 0.193 | -0.082 | 0.434 | 0.317 | 0.373 | 0.073 | -0.003 |
|  | P-value | 0.221 | 0.604 | 0.004 | 0.041 | 0.015 | 0.645 | 0.987 |
| RSN | Cor | -0.020 | -0.08 | 0.216 | 0.350 | 0.201 | 0.158 | 0.064 |
|  | P value | 0.714 | 0.143 | < 0.001 | < 0.001 | < 0.001 | 0.004 | 0.245 |

**Table 4.** Results of Spearman’s correlation test corresponding to Figure 4B.

| Period |  | Start1 | Start2 | Cue1 | Cue2 | Cue3 | Cue4 | prefb |
| --- | --- | --- | --- | --- | --- | --- | --- | --- |
| FSN | rho | 0.213 | -0.019 | 0.359 | 0.385 | 0.373 | -0.017 | -0.076 |
|  | P value | 0.175 | 0.906 | 0.020 | 0.012 | 0.015 | 0.917 | 0.632 |
| RSN | rho | -0.011 | -0.070 | 0.162 | 0.288 | 0.145 | 0.131 | 0.054 |
|  | P value | 0.836 | 0.200 | 0.003 | < 0.001 | 0.008 | 0.016 | 0.322 |

In conclusion of the z-scored firing-rate analysis, these similarities indicated that the extent of neural modulations (i.e., response sensitivity in raw firing rates and signal contribution in z-scored firing rates) was greater in FSNs than that in RSNs, suggesting that the stronger neural modulation observed in FSNs was not due to the inflation of the estimated regression coefficients by the noise, which is sometimes suspected to occurr in the higher raw firing rates analysis. Namely, encoded information for probability and magnitude (i.e., signal) must be stronger in FSNs than those in RSNs (Figure 4C).

### Dynamic range and information processing during expected value computation: Analyses in the raw firing rates

Next, we examined how expected value computation relies on the basic firing properties of FSNs and RSNs (Figure 2) by analyzing the influence of dynamic range on the extent of neural modulation, a key factor regulating cortical computation according to the local circuit structure (Kawaguchi, 1995). In this analysis, we also applied the same analyses of the regression coefficients on both raw and z-scored firing rates to compare the results and keep consistency of our analyses. First, we describe the results based on the raw firing rates.

When we analyzed the regression coefficients for the raw firing rates, we found that the dynamic range affected the extent of neural modulation in both FSNs and RSNs, that is, the regression coefficients showed dynamic range dependency (Figure 5A, general linear model, n = 3770 [377 neurons ₓ 5 task periods ₓ 2 coefficient types], dynamic range: *F* = 1112.7, *P* < 0.001, df = 1). Unambiguously, a wider dynamic range in FSNs co-occurred with stronger neural modulation in FSNs (Figure 5A, neuron type: *F* = 32.3, *P* < 0.001, df = 1), whereas there was no significant difference between the probability and magnitude modulation types (Figure 5A, green and blue, coefficient type: *F* = 2.08, *P* = 0.150, df = 1). Thus, both FSNs and RSNs process expected value computations under the influence of dynamic range in raw firing rates, though stronger dependency on the dynamic range can be possibly explained by both signal and noise.

**Figure 5.**
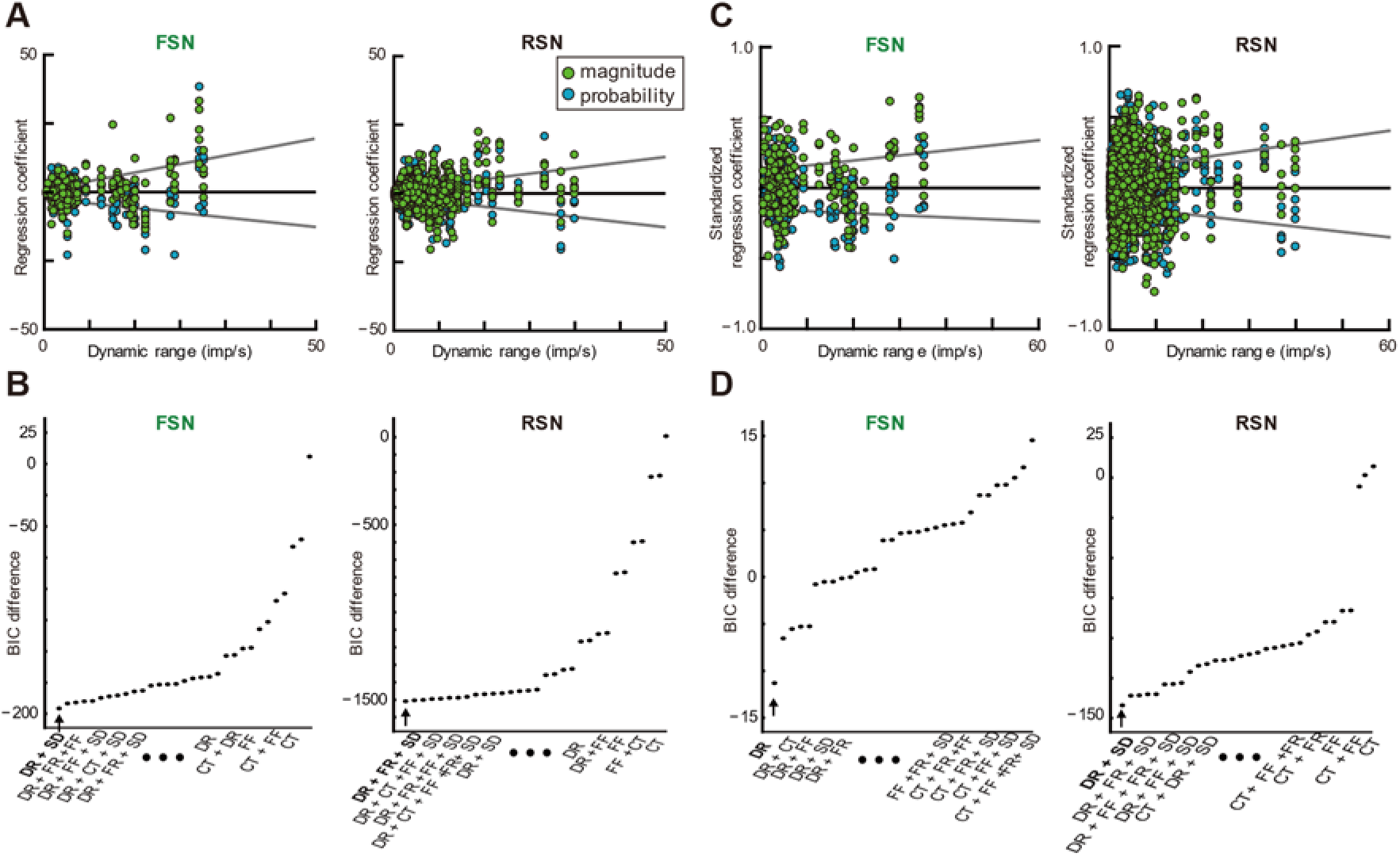
Dynamic range of firing rates differed between FSNs and RSNs in neural modulation. **A.** Plots of regression coefficients for the probability (blue) and magnitude (green) of rewards against the dynamic range for FSNs (left) and RSNs (right). The gray lines represent the regression slopes from the general linear model. **B.** Differences in Bayesian information criterion (BIC) values between the models and the null model. The x-axis labels indicate the selected models in rank order. Names of top and worst five models are shown: DR, dynamic range; FR, firing rate; CT, regression coefficient type (i.e., probability or magnitude); FF, fano factor; SD, standard deviation of firing rates. The arrowheads indicate the BIC values for the best models. Bold indicates the best model. **C–D.** Same as A–B, but for analyses using z-scored firing rates (i.e. standardized regression coefficient).

We also applied a model selection approach to explore the factors that best explained the neural modulations in FSNs and RSNs. We found that the combination of the standard deviation of the firing rates in each task period (SD) and dynamic range (DR) best explained information processing in FSNs (Figure 5B; best model in bold; log-likelihood ratio test, *P* < 0.001 for all conditions). The similar model best explained the extent of neural modulation in RSNs, DR + SD + FR (mean firing rate). Thus, FSNs and RSNs might share some information processing in the OFC circuit during expected value computation.

### Dynamic range and information processing during expected value computation: Analyses of z-scored firing rates

To explore the mechanisms for expected value computation, we repeated the analyses using regression coefficients for the z-scored firing rates, i.e., standardized regression coefficients estimated in each task period. The results were slightly different from those of the raw firing rates, as follows: The models best explaining the differences in neural modulations differed between FSNs and RSNs (Figure 5D, see the x-axis label for the selected models in rank order). The DR-only model best explained the FSNs’ z-scored firing rates, in contrast to the RSNs, whose best model was DR + SD, indicating dynamic range is a key factor in the encoded signal of OFC circuit.

Additionally, while neural modulations of both FSNs and RSNs showed dynamic range dependency (Figure 5C, general linear model, n = 3770 [377 neurons ₓ 5 task periods ₓ 2 coefficient types], dynamic range: *F* = 143.3, *P* < 0.001, df = 1), the extent of dependency did not differ between FSNs and RSNs (Figure 5C). Unambiguously, there was no significant difference between FSNs and RSNs for dynamic range dependency of the regression coefficients in the z-scored firing rates (Figure 5C, neuron type: *F* = 0.41, *P* = 0.52, df = 1, coefficient type: *F* = 1.82, *P* = 0.178, df = 1), in contrast to the results of the raw firing rates (Figure 5A, neuron type: *F* = 32.1, *P* < 0.001, df = 1, coefficient type: *F* = 2.07, *P* = 0.15, df = 1). The z-scored firing rate results suggested that the encoded information in both FSNs and RSNs similarly depended on the dynamic range during expected value computation.

Collectively, when we analyzed the z-scored firing rates, the neural modulations were similarly dependent on the dynamic range in both RSNs and FSNs (Figure 5C and D), in which the influence of noise was removed and the encoded information (i.e., signal) contributed to this observation. Because z-scored firing rates allow us to fairly compare encoded information between two neural populations, these results indicate that encoded information in FSNs and RSNs may be similarly controlled by the dynamic rage (Figure 5C), but stronger signal appeared in FSNs during expected value computations (Figure 4C).

## DISCUSSION

In the present study, we analyzed the activity of OFC neurons recorded during economic behaviors in monkeys. We differentiated FSNs from other neurons (i.e., RSNs) based on spike waveforms. Thereafter, we identified two properties inherent to FSNs relative to RSNs. First, FSNs displayed high-frequency firing rates, high across-trial variance, wide dynamic ranges, and more regular firing intervals than RSNs. Second, the neural representation of the probability and magnitude of rewards (i.e., neural modulations for economic behavior) was similar but quantitatively different between the two classes, with FSNs encoding reward information in a manner similar to RSNs in terms of the proportion of neurons (Figure 4A). In addition, both FSNs and RSNs represent expected values at the population level (Figure 4B, Cue1 period), but the extent of neural modulations in FSNs was larger than those in RSNs (Figure 4C), Furthermore, we found that the dynamic range is a key factor in explaining these neural modulations in both FSNs and RSNs, with similar dependences on the dynamic range in FSNs and RSNs (Figure 5C). Since the z-scored firing rates allow direct comparison of encoded signals between two neural populations, these results suggest that i) FSNs have stronger signals than RSNs in expected value computation, and ii) FSNs and RSNs may share information processing in encoding expected value in monkey OFCs.

### Identification of FSNs in the primate OFC

In vivo, FSNs have been identified based on extracellularly recorded spike waveforms in rodents (Bartho et al., 2004). Bartho et al. identified neurons in the rat prefrontal cortex based on a narrow spike waveform recorded extracellularly, which reflected the intracellular properties of action potentials (Henze et al., 2000). Most of the identified neurons showed inhibitory effects on neighboring neurons, whereas none showed excitatory effects (Figure 4 in Bartho et al., 2004), indicating that these narrow spike-waveform neurons seemed to be inhibitory interneurons. Accumulating evidence from in vivo and in vitro studies of cortical and subcortical structures supports the hypothesis that narrow spike-waveform neurons are parvalbumin-containing GABAergic interneurons (FSNs) in rodents (Chamberland et al., 2023; Gage et al., 2010; Giordano et al., 2023; Hatch et al., 2017; Inokawa et al., 2010; Owen et al., 2018) and in monkeys (Gonzalez-Burgos et al., 2005; Kunimatsu et al., 2021; Yamada et al., 2016). The electrophysiological and neurochemical properties of cortical and subcortical structures are similar in primates and rodents (Kawaguchi and Kondo, 2002; Kawaguchi et al., 1995), and it is generally agreed that FSNs recorded from behaving monkeys are parvalbumin-containing GABAergic interneurons.

In the present study, we identified FSNs based mainly on spike waveforms (Figure 2A, green), similar to previous rodent studies of cortical and subcortical structures (Bartho et al., 2004; Gage et al., 2010). The identified FSNs exhibited high-frequency firing rates during the task period (approximately 10 Hz) compared to RSNs (Figure 2C). However, the average firing rate of the FSNs in this study was lower than that reported in other monkey studies in the visual cortex (>30 Hz) (Mitchell et al., 2007) and prefrontal cortex (>20 Hz) (Fan et al., 2017). This discrepancy may arise from differences in cortical regions as well as behavioral tasks performed by the monkeys, because neural firing rates depend on input to the local circuit (Figure 6, gray), although cortical areas share a six-layer structure composed of different types of interneurons (Kawaguchi and Kondo, 2002).

**Figure 6.**
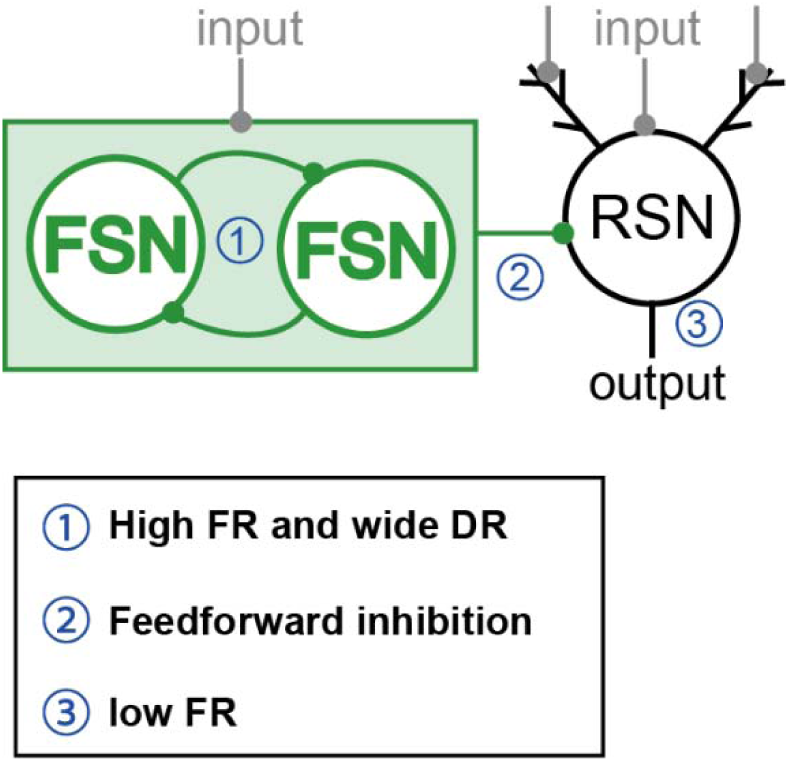
Schematic depiction of FSN-RSN neural circuit for information processing. Schematic anatomical depiction of the cortical circuit for the presumed parvalbumin-containing GABAergic interneurons (FSNs) and presumed output pyramidal neurons (RSNs) (top). This figure has been modified from Figure 1a, by Hattori et al. (2017).

The spike waveform is one of the predominant characteristics used to identify neuron types in vivo. However, it cannot differentiate between all neuron types. RSNs appeared to comprise multiple neuron types with differences in spike irregularity (Figure 2M). In addition, spike waveforms are strongly dependent on the amplifier filter settings: the frequency of the low-pass and high-pass filters and the type of filter (e.g., Butterworth, Bessel, or Chebyshev) (Yamada et al., 2016); hence, the characteristics of the spike waveform must be compared under the same experimental settings. Thus, we reliably identified FSNs in the present study.

We note that the majority of the FSNs and RSNs in this study was recorded from deeper OFC layers, such as four, five and six, because we recorded neurons just after we found OFC cortex during penetration of electrode (In OFC six layer to first layer appeared sequentially through top to bottom during a penetration of electrode).

### Dynamic range, firing rates, and neural modulations in the raw firing rates

In the present study, we found similar but slightly stronger neural modulations for probability and magnitude in FSNs than in RSNs at the neuronal population level (Figure 3E–F) in the analysis of raw firing rates. This finding contrasts with that observed in the striatum, where FSNs are less selective than output neurons (Yamada et al., 2016). While local circuit structures differ between cortical and subcortical regions (Inokawa et al., 2020; Isomura et al., 2009; Tsubo et al., 2013), FSNs in both consistently showed high-frequency baseline firing rates that inhibited neighboring neurons. The higher firing rates in FSNs may result from intrinsic membrane properties, such as high input resistance (Kawaguchi, 1993, 1995), although other possibilities also exist, including differences in morphology, synapse density, and other cellular properties. If neurons can easily become active in response to excitatory inputs, high-frequency firing rates (Figure 2C, L) and larger changes in task-related activity should occur; hence, a wider dynamic range (Figure 2K) should be observed. These neural properties are related to larger changes in neural modulation as a function of firing rate and dynamic range (Figure 5A, compare FSNs and RSNs’ regression slopes). Consequently, the output neurons in the cortical (Chen et al., 2023; Imaizumi et al., 2022; Yamada et al., 2021; Yamada et al., 2018) and subcortical (Chen et al., 2025; Yamada et al., 2013; Yamada et al., 2011; Yamada et al., 2007) structures are inhibited via feedforward inhibition (Figure 6) during economic behavior.

We caution that these stronger neural modulations observed in the raw firing rates likely reflect increases in both signal and noise; therefore, z-scored firing rates are needed to compare the encoded signals of FSNs and RSNs. This issue is discussed further in the following sections.

### Influences of the dynamic range between the raw and z-scored firing rates

We observed a similar dynamic-range dependency on neural modulation between FSNs and RSNs when the z-scored firing rates were analyzed (Figure 5C), in contrast to the result from the raw firing rate analysis (Figure 5A). What does this opposite result indicate? First, neural modulations, as observed in the regression coefficients, reflect both signal and noise when raw firing rates are analyzed. However, considering that information processing occurs within the circuitry, how much signals are encoded by the neurons is important. We compared the encoded information between FSNs and RSNs using z-scored firing rates (Chung et al., 1987; Passaglia and Troy, 2004; Theunissen et al., 1996), and found stronger neural modulations in FSNs (Figure 4C). However, the encoded information was similarly dependent on dynamic range in FSNs and RSNs (Figure 5C). These results suggest that shared information processing between these two types of neurons may occur under the influence of dynamic range in the OFC circuit. To clarify these circuit mechanisms, further studies are required to examine neuronal activity by directly labeling and observing the activity of connected neuron pairs.

We found that the relationships in reward processing for probability and magnitude were similar but slightly different between FSNs and RSNs (Figures 3–5), irrespective of whether raw or z-scored firing rates were used. These similarities and differences between FSNs and RSNs may arise from the local circuit structure: mutual inhibition between FSNs and feedforward inhibition from FSNs to RSNs (Figure 6, green). Mutual inhibition determines the mean firing rates of circuitry neurons according to the excitatory input level, whereas feedforward inhibition determines the output level of the circuit, namely, RSN activity. Although excitatory inputs were not observed in this study, these two key properties of the local circuit can regulate the expected value computations. Indeed, recurrent inhibition controls the circuit dynamics (Lynn et al., 2025).

## Limitations of the study

One limitation of our study is that the functional difference of FSNs between cOFC and mOFC is not obvious in this study. While we have previously reported the weaker neural modulation in mOFC than that in cOFC at the population level (Chen et al., 2023; Chen et al., 2025; Imaizumi et al., 2022; Yamada et al., 2021), we were not able to surely compare the role of FSNs between these regions due to the small number of samples (cOFC, 25; mOFC, 17).

Another limitation of our study is that we did not examine the activity of directly connected FSN–RSN pairs. Therefore, we could not test any hypothesis for the information processing based on the local circuit connection, directly. All our testing of computational processes was supported at the population level, but not for the FSN–RSN pair. While direct examination of connected FSN–RSN pairs was not performed in the current study or most previous studies in the prefrontal cortex, striatum, and hippocampus, no study has identified the activity of FSNs in the monkey OFC during economic behavior.

### Summary: coding of reward probability and magnitude information by FSNs and RSNs

Our results for the z-scored firing rates suggest that FSNs may share information processing with RSNs in the OFC, showing a stronger sensitivity to probability and magnitude (Figure 4B and C). These encoded signals were similarly influenced by the dynamic range in both FSNs and RSNs (Figure 5C). This observation is consistent with previous literature. For example, inhibitory interneurons have been suggested to play a functional role in regulating neural signals in the cerebral cortex. FSNs with parvalbumin immunoreactivity in the visual area V1 of mice have been shown to be selectively involved in shaping orientation tuning and enhancing the directional sensitivity of neighboring neurons (Lee et al., 2012). Furthermore, FSNs have been suggested to play an inhibitory role in improving various cognitive functions in distinct cortical regions. For example, FSNs in the monkey prefrontal cortex are associated with learning and performance of cognitive tasks (Constantinidis and Goldman-Rakic, 2002; Qi and Constantinidis, 2012). FSNs in the visual area V4 show modulation in their control of attention (Mitchell et al., 2007), suggesting that the reliability of the output neuron response is increased by reducing response variability. Previous monkey studies on other prefrontal regions have also indicated that FSN activity is selective for reward cues, with some similarity to RSNs (Fan et al., 2017; Kawai et al., 2019). In our study, stronger signal carried by FSNs may shape RSNs signals (i.e., output) dependent on firing rate level. Feedforward inhibition must be a general mechanism for improving output sensitivity, whereas the input structure is the key factor in driving a local network.

## Materials and Methods

### Subjects and experimental procedures

Two rhesus monkeys were used in this study (*Macaca mulatta,* SUN, 7.1 kg, male; *Macaca fuscata*, FU, 6.7 kg, female). All experimental procedures were approved by the Animal Care and Use Committee of the University of Tsukuba (protocol no. 23-057) and performed in compliance with the U.S. Public Health Service’s Guide for the Care and Use of Laboratory Animals. Each animal was implanted with a head-restraining prosthesis. Eye movements were measured using a video camera at 120 Hz. Visual stimuli were generated using a liquid-crystal display at 60 Hz, placed 38 cm from the monkey’s face when seated. The subjects performed the cued lottery task five days a week. The subjects practiced the cued lottery task for ten months, after which they became proficient in choosing lottery options. We have previously reported the activity of RSNs, but not that of FSNs, during this task.

### Behavioral task

#### Cued lottery tasks

The animals performed one of two visually cued lottery tasks: a *single cue task* or a *choice task*. Neuronal activity was recorded only during the single-cue task. Single-cue task: At the beginning of each trial, the monkeys had 2 s to align their gaze within 3^°^ of a 1^°^-diameter gray central fixation target. After fixing for 1 s, an 8^°^ pie chart providing information on the probability and magnitude of the rewards was presented for 2.5 s at the same location as the central fixation target. The pie chart was then removed, and 0.2 s later, a 1 kHz or 0.1 kHz tone of 0.15 s duration indicated the reward and no-reward outcomes, respectively. The animals received a fluid reward, with the magnitude and probability indicated by green and blue pie charts, respectively; otherwise, no reward was delivered. A high tone preceded the reward by 0.2 s. A low tone indicated that no reward was delivered. An intertrial interval of 4–6 s followed each trial.

Choice task: At the beginning of each trial, the monkeys had 2 s to align their gaze within 3^°^ of a 1^°^-diameter gray central fixation target. After fixing for 1 s, two peripheral 8^°^ pie charts providing information on the probability and magnitude of the rewards for each of the two target options were presented for 2.5 s, at 8^°^ to the left and right of the central fixation location. The gray 1°choice targets appeared at the same location. After a 0.5 s delay, the fixation target disappeared, cueing saccade initiation. The animals were free to choose for 2 s by shifting their gaze to either target within 3^°^ of the choice target. A 1 kHz or 0.1 kHz tone of 0.15 s duration indicated reward and no-reward outcomes, respectively (i.e., feedback sounds). The animals received a fluid reward, indicated by the green pie chart of the chosen target, with the probability indicated by the blue pie chart. Otherwise, no reward was delivered. An intertrial interval of 4–6 s followed each trial.

#### Pay-off and block structure

Green and blue pie charts indicated reward magnitudes from 0.1 to 1.0 mL, in 0.1 mL increments, and reward probabilities from 0.1 to 1.0, in 0.1 increments, respectively. One hundred pie charts were used in this study. In the single-cue task, each pie chart was presented once in a random order. In the choice task, two pie charts were randomly assigned to the two options. During one session of electrophysiological recording, approximately 30–60 trial blocks of the choice task were sometimes interleaved with 100–120 trial blocks of the single-cue task.

#### Calibration of the reward supply system

The precise amount of liquid reward was controlled and delivered to the monkeys using a solenoid valve. An 18-gauge tube (0.9 mm inner diameter) was attached to the tip of the delivery tube to reduce variation across trials. The reward amount for each payoff condition was calibrated by measuring the weight of water with a precision of 0.002 g (2 μL) on a single-trial basis. The calibration method was the same as described previously (Yamada et al., 2018).

### Electrophysiological recordings

Conventional techniques were used to record single-neuron activity in the cOFC and mOFC. Monkeys were implanted with recording chambers (28 × 32 mm) targeting the OFC and striatum, centered 28 mm anterior to the stereotaxic origin. The locations of the chambers were verified using anatomical magnetic resonance imaging (MRI). At the beginning of each daily recording session, a stainless-steel guide tube was placed within a 1-mm spacing grid, and a tungsten microelectrode (1-3 MΩ, FHC) was passed through the guide tube. To record the neurons in the mOFC and cOFC, the electrode was lowered until it reached the bottom of the brain after passing through the cingulate cortex, dorsolateral prefrontal cortex, or between them. Electrophysiological signals were amplified, bandpass filtered, and monitored. Single-neuron activity was isolated based on the spike waveforms. We recorded from the two brain regions of a single hemisphere of each monkey (179 neurons in monkey SUN and 198 in monkey FU): 42 FSNs (cOFC, 25; mOFC, 17) and 335 RSNs (cOFC, 182; mOFC, 153). The activities of each neuron were sampled only when it demonstrated a good signal-to-noise ratio (>2.5). No blinding was conducted. The sample sizes required to detect the effect sizes (number of recorded neurons, number of recorded trials in a single neuron, and number of monkeys) were estimated according to previous studies (Enomoto et al., 2020; Yamada et al., 2013; Yamada et al., 2018; Yamada et al., 2004). Neural activity was recorded during 100–120 trials of the single-cue task. During the choice trials, neural activity was not recorded.

### Classification of neuron type

In this analysis, FSNs (presumed parvalbumin-containing GABAergic interneurons) were differentiated from RSNs (presumed pyramidal neurons) based on their spike width (i.e., the width at half maximum of the negative peak amplitude and the width from peak to valley), according to a previous study (Bartho et al., 2004). In addition, the coefficient of variation of the inter-spike intervals (Maimon and Assad, 2009) was used. Clustering of spike waveforms based on principal component analysis was performed using the prcomp() function in R. A dendrogram was constructed to verify the clusters. FSNs were classified as neurons in the cluster exhibiting narrow spike waveforms, whereas all other neurons were classified as RSNs. In our previous reports that used this dataset (Chen et al., 2023; Chen et al., 2025; Imaizumi et al., 2022; Yamada et al., 2021), we reported the activity of RSNs, but not of FSNs. The number of RSNs reported in this study differs from previous studies because we did not perform a quantitative classification of neuron types.

## Statistical analysis

Statistical analyses were performed using the R statistical software package (R Foundation for Statistical Computing, Vienna, Austria, http://www.r-project.org/). All statistical tests for behavioral and neural analyses were two-tailed.

### Effects of units on statistical analysis

In this study, we used two variables for analysis: probability and magnitude. We defined reward probability from 0.1 to 1.0 and reward magnitude from 0.1 to 1.0 mL. Under this definition, the effects of probability and magnitude on the data were equivalent.

### Behavioral analysis

No new behavioral results were included; however, the procedure for behavioral analysis was as follows: We previously reported that monkey behavior depends on expected values, defined as the probability multiplied by the magnitude (Yamada et al., 2021). We described the analysis steps to check whether the monkey’s behavior reflected task parameters such as reward probability and magnitude. Importantly, we showed that the monkeys’ choice behavior reflected the expected values of the rewards, that is, the probability multiplied by the magnitude. For this purpose, the percentage of trials in which the right option was chosen was analyzed in the pooled data using a general linear model with a binomial distribution:

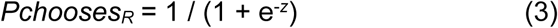

where the relationship between *Pchooses_R_* and *Z* is given by a logistic function in each of the following three models: number of pie segments (M1), probability and magnitude (M2), and expected values (M3).

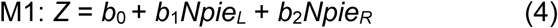

where *b*_0_ is the intercept and *Npie_L_* and *Npie_R_* are the number of pie segments contained in the left and right pie chart stimuli, respectively. The values of *b*_0_ to *b*_2_ are free parameters and are estimated by maximizing the log likelihood.

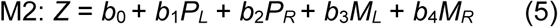

where *b*_0_ is the intercept; *P_L_* and *P_R_* are the probabilities of rewards for the left and right pie chart stimuli, respectively; and *M_L_* and *M_R_* are the magnitudes of the rewards for the left and right pie chart stimuli, respectively. The values of *b*_0_–*b*_4_ are free parameters and were estimated by maximizing the log likelihood.

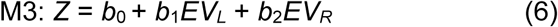

where *b*_0_ is the intercept and *EV_L_* and *EV_R_* are the expected values of rewards as probability multiplied by magnitude for the left and right pie chart stimuli, respectively. The values of *b*_0_ to *b*_2_ are free parameters and are estimated by maximizing the log likelihood. We identified the best model describing monkey behavior by comparing goodness-of-fit using Akaike’s information criterion (AIC) and the Bayesian information criterion (BIC) (Burnham and Anderson, 2004).

### Neural analysis

Peristimulus time histograms were drawn for each neuron’s activity, aligned at the onset of a visual cue. The average activity curves were smoothed using a 50-ms Gaussian kernel (σ = 50 ms). We analyzed neural activity during the 2.5-s pie chart stimulus presentation in the single-cue task, including baseline activity during the 1.0-s fixation period before cue presentation. The firing rates of each neuron during the 0.5-s time window were estimated every 0.5 s for a total of seven analysis periods: Start1, Start2, Cue1, Cue2, Cue3, Cue4, and Pre-fb (feedback). A Gaussian kernel was not used for the statistical analyses.

Basic firing properties—peak firing rates, peak latency, duration of peak activity (half-peak width), dynamic range (DR), and coefficient of variation of the inter-spike interval (CV ISI)—were compared between FSNs and RSNs using parametric or nonparametric tests at a significance level of *P* < 0.05. The dynamic range was defined as the difference between the maximum and minimum firing rate across the seven task periods after the start of fixation: Start1, Start2, Cue1, Cue2, Cue3, Cue4, and Pre-fb. Baseline firing rates 1 s before the appearance of the central fixation targets were also compared at a significance level of *P* < 0.05. We also estimated variability of spikes during the seven analysis periods, such as i) across-trial variance estimated as S.D. and ii) fano factor estimated as S.D. divided by mean. Yamada et al. (2021) also analyzed the some of these basic firing properties of RSNs, but not FSNs.

#### Z-scored and raw firing rates

For the subsequent analyses, we used both raw and z-scored firing rates. The z-scored firing rates were computed by standardized the firing rates in each analysis period.

*Linear regression used to detect firing modulations in each neuron’s activity.* Neural discharge rates (*F*) were fitted using the following variables:

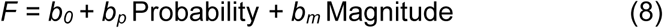

Where Probability and Magnitude are the probability and magnitude of the rewards indicated by the pie chart, respectively. *b*_0_ is the intercept. If *b_p_* and *b_m_* were not zero at *P* < 0.05, the discharge rates were considered significantly modulated by the corresponding variable.

On the basis of the linear regression, activity modulation patterns were categorized into several types: “Probability” (P) type with a significant *b*_p_ and without a significant *b*_m_; “Magnitude” (M) type without a significant *b*_p_ and with a significant *b*_m_; “Expected value” (EV) type with significant *b*_p_ and *b*_m_ with the same sign (i.e., positive *b*_p_ and positive *b*_m_ or negative *b*_p_ and negative *b*_m_); “Risk-Return” (RR) type with significant *b*_p_ and *b*_m_ with both having opposite signs (i.e., negative *b*_p_ and positive *b*_m_ or positive *b*_p_ and negative *b*_m_) and “non-modulated” type without significant *b*_p_ and *b*_m_. The risk–return types reflect high-risk high returns (prefer low probability and large magnitude) or low-risk low returns (prefer high probability and low magnitude).

We compared the basic firing properties and activity modulations between FSNs and RSNs as follows: 1) proportion of neuron types using the chi-square or multinomial test; 2) average firing rates using ANOVA, Kruskal-Wallis test, or Wilcoxon rank-sum test with Bonferroni correction for multiple comparisons; and 3) regression coefficients using a general linear model, such as ANOVA and linear regression.

#### Analysis of z-scored firing rates in each neuron’s activity

To fairly compare neuronal activity modulations between FSNs and RSNs, we estimated the z-scored firing rates. We normalized each neuron’s activity to z-scores in each analysis period (firing rates during inter-trial intervals), as mentioned above. Thereafter, we performed the same analyses described above for *F* in Eq. 8 using the z-scored firing rates. We estimated standardized regression coefficient in each neural activity during each analysis period.

*Linear regression to detect firing modulations at the population level.* The regression coefficients for reward magnitude (*R*) were fitted using the following variables:

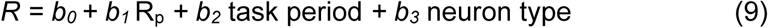

where R_p_ denotes the regression coefficient of reward probability. The task period was a categorical variable consisting of Cue1, Cue2, Cue3, Cue4, and Pre-FB. The neuron type was a categorical variable comprising FSNs and RSNs. If *b_1_*to *b_3_* were not zero at *P* < 0.05, the regression coefficients were considered significantly modulated by the corresponding variable.

#### Dynamic range and neural modulations

To analyze the influence of basic firing properties on the regression coefficients for the probability and magnitude of rewards, we modeled how the dynamic range and average firing rates affected neural modulation as follows:

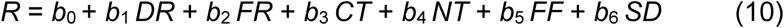

Where *R* is the absolute value of the regression coefficients for reward probability and magnitude (b_p_ and b_m_) in Eq. 8, *b*_0_ is the intercept. *DR* is the dynamic range. *FR* is the average firing rate in each of the five task periods after cue presentation: Cue1, Cue2, Cue3, Cue4, and Pre-FB. *CT* is the regression coefficient type (i.e., probability or magnitude) as a categorical parameter. *NT* is a categorical variable representing neuron type: FSN or RSN. *FF* is fano factor. *SD* is the standard deviation of firing rates across trials. If *b_1_* was not 0 at *P* < 0.05, neural modulation by reward probability and magnitude was considered to be significantly influenced by the dynamic range of the neurons. If *b_2_* was not 0 at *P* < 0.05, neural modulation by reward probability and magnitude was considered to be significantly influenced by each neuron’s average firing rate. If *b_3_*was not 0 at *P* < 0.05, neural modulation differs between reward probability and magnitude, that is, between the regression coefficient types. If *b_4_* was not 0 at *P* < 0.05, neural modulation by reward probability and magnitude differed between FSNs and RSNs. If *b_5_* was not 0 at *P* < 0.05, neural modulation by reward probability and magnitude was considered to be significantly influenced by the fano factor. If *b_6_*was not 0 at *P* < 0.05, neural modulation by reward probability and magnitude was considered to be significantly influenced by the standard deviation of firing rates. We evaluated the results using both raw and z-scored firing rates.

The regression coefficient values were also compared using ANOVA, including neuron type (FSN or RSN), coefficient type (probability or magnitude), coding sign (positive or negative value), and task period (Cue1 to pre-feedback periods). For this analysis, the absolute value of the regression coefficient was used to detect the significant main effects of the four factors at P < 0.05. Regression coefficients from both raw and z-scored firing rates were used in this analysis.

#### Model comparisons

To identify the best structural model for neural modulation, we applied a model-selection approach using all possible combinations of variables in Eq. 10, as described above. We sought a combination of best-fit parameters to explain the neural modulation based on the probability and magnitude of rewards. We compared the goodness of fit using BIC.

After estimating the best-fit parameters for each model, the model with the lowest BIC values was selected. To evaluate model fit, we calculated the difference between these values and the null model’s BIC, which is the log likelihood under the assumption that all free parameters are zero in the model, except for the intercept *b*_0_. We used the log-likelihood ratio test for each of the selected null models at *P* < 0.05.

## Acknowledgements

The authors express their appreciation to Kaoru Kouguchi, and Shiho Nishino for technical assistance. We appreciate Yasuhiro Tsubo for his valuable comments. Monkey FU was provided by NBRP “Japanese Monkeys” through the National Bio Resource Project of MEXT, Japan. This study was supported by JSPS KAKENHI grant numbers JP15H05374 and JST Moonshot R&D JPMJMS2294 (H.Y.).

## Author Contributions

H.Y. designed the study; Y.I. and H.Y. conducted the experiments; T.M., T.K., and H.Y. analyzed the data; and H.Y. and T.K. wrote the manuscript. All authors approved the final manuscript.

## Data Availability

All data used in this study are presented in the manuscript.

